# AAV gene therapy for Cockayne syndrome

**DOI:** 10.1101/2025.06.06.658349

**Authors:** Ana Rita Batista, Aine C. Scholand, William S. Callahan, McKenna K. Watson, Cassandra M. Sion, Tyler Mola, Kennedy O’Hara, Oliver D. King, Robert M. King, Miguel Sena-Esteves

## Abstract

Cockayne Syndrome (CS) is an autosomal recessive, progressive developmental and neurodegenerative disease. Approximately 30% of cases are caused by mutations in the *ERCC8/CSA* gene. Patients with CS present with cutaneous photosensitivity, growth failure, shorter life span and a progressive degeneration of the central nervous system. Loss of function mutations in CSA result in deficiencies in transcription-coupled nucleotide excision repair, regulation of RNA Pol II mediated transcription repair of oxidative DNA damage, and mitochondrial metabolism. Currently there are no available therapies for these patients. AAV gene therapy offers an opportunity to address this unmet need. We designed a new AAV vector encoding human CSA under a CBA promoter. We tested the therapeutic efficacy of this AAV9-CSA vector by neonatal ICV injection in the *Csa^-/-^;Xpa^-/-^* mouse model. Treatment with AAV9-CSA resulted in a significant increase in lifespan, and broad distribution of human CSA in the brain and heart. Despite clear therapeutic benefit, we also observed neuroradiological abnormalities, neuropathologic alterations including hypo-myelination, astrocytosis, microgliosis, and likely life limiting transcriptomic alterations in liver at endpoint. Nonetheless, the success of these experiments paves the way for the first in human clinical translation of a gene therapy for CS patients.

## INTRODUCTION

Cockayne Syndrome (CS) is a multi-system, autosomal recessive, progressive developmental and neurodegenerative disorder, caused primarily by mutations in either the *ERCC8* (Cockayne Syndrome A or *CSA*) or *ERCC6* (Cockayne Syndrome B or *CSB*) gene. It is very rare, devastating and almost universally fatal in children. The incidence of CS is 1 per 250,000 live births, but it is frequently undiagnosed (1-4). The fundamental signs of disease are microcephaly, growth failure, and developmental delay. Other common features include short stature, cataracts, sunken eyes, hearing loss, cutaneous photosensitivity, liver dysfunction, dental caries and morphological abnormalities of the teeth (5-8). The central nervous system (CNS) is the most heavily impacted by CS and its pathology contributes significantly to morbidity, although the most common cause of death is pneumonia and recurrent respiratory infection (6, 9).

*ERCC8 (CSA)* and *ERCC6* (*CSB)* mutations account for approximately 30% and 70% of cases of CS, respectively (6). Mutations in these genes result in defects in transcription-coupled nucleotide excision repair (TC-NER) leading to gene misregulation and cytotoxicity due to RNA polymerase II stalling in non-dividing tissues (10). Additionally, CSA and CSB proteins are involved in repair of oxidative DNA damage (11, 12), and mitochondrial metabolism (13), which contribute to the growth failure and neurodegeneration seen in children. Our research focuses on the rarest form of CS caused by mutations in the *ERCC8/CSA* gene.

CSA is a 44 kDa protein that contains seven WD40 repeats (14). CSA was originally described as a key player in the TC-NER pathway (14-16), which is responsible for the repair of bulky DNA lesions, such as those caused by UV light and/or redox damage. The triggering TC-NER event is the arrest of RNA polymerase II (RNA pol II) due to the presence of lesions in the actively transcribed strand of a gene. CSA is part of the E3 ubiquitin ligase complex (17, 18) responsible for ubiquitination and degradation of TC-NER proteins once repair is complete, which is required for RNA pol II to re-start transcription (15, 19-21). In the absence of CSA protein, the TC-NER complex normally composed of CSA, CSB, XPA, etc, remains attached to the DNA and the arrest of RNA pol II persists, leading to p53 activation and cell death. However, TC-NER defects alone do not explain the severe neurological disease observed in CS (22, 23). Besides proteins in the TC-NER complex, CSA also interacts with several other proteins involved in transcription, ribosomal biogenesis, and mitochondrial homeostasis, but the exact molecular mechanisms are not fully understood. All of these are cellular and molecular phenotypes of CS (24).

Several animal models of CS have been developed over the years to understand the disease mechanism. The first *Csa^-/-^* mouse model only exhibits pronounced skin cancer after chronic exposure to UV light and marked photoreceptor loss, but otherwise has no other apparent phenotypes or pathological alterations and has a normal lifespan (25). This mouse model also fails to reproduce the myriad of somatic and neurologic deficits of CS, with the exception of clusters of activated microglia observed in close association with mature oligodendrocytes, which resemble the patchy demyelination observed in patients (22, 26). To generate a more aggressive CS animal model, a double knockout CX mouse was generated by deleting *CSA* as well as *Xpa,* which encodes another protein part of the TC-NER complex. Although this is a genetically artificial CS model, it is a strong surrogate for testing new therapies, as it faithfully recapitulates the neurological features of CS. The CX model is indistinguishable from controls at birth but has impeded growth by postnatal day 5. By day 12 CS disease is evident, with a median survival of 22 days. Signs of CS disease include kyphosis, abnormal gait consistent with ataxia, uncoordinated hind limb movement, dystonia, and eventual paralysis (27).

There are no disease-modifying treatments to halt or slow CS disease progression. Current treatment consists of supportive care and interventions to manage the disease manifestations. Since CS is a monogenic disease, with loss-of-function mutations, it is amenable to gene replacement therapy. AAV vectors are a powerful and promising tool for delivery of gene therapies, with several therapies already approved by the FDA, including CNS applications, and with a considerably number of ongoing clinical trials (28). Here we are the first to develop an AAV9 gene therapy for Cockayne Syndrome showing dramatic phenotypic rescue using an aggressive CS mouse model.

## RESULTS

To test the efficacy of our new scAAV9-CSA vector *in vitro*, we generated a new cell line of HEK293T-CSA^-/-^ cells (see Supplemental Fig.1). Upon infection of CSA ^-/-^ cells with AAV9-CSA vector (1E+5 vg/cell), CSA protein expression was equivalent to normal levels in HEK293T cells (Fig. 1A). When using fibroblasts derived from CS patients transduced with an AAV3b-CSA vector (dose of 1E+5 vg/cell), CSA protein expression was higher than in normal human fibroblasts (Fig. 1B). We also used a functional assay to demonstrate the biological activity of the CSA protein by exposing cells to the chemotherapeutic drug Illudin S, which is a potent DNA alkylating agent. DNA lesions caused by this genotoxic drug are repaired exclusively by TC-NER (29), causing CSA-deficient cells to die (20). In our assays, transduction with AAV-CSA vectors promoted survival of both HEK293T-CSA^-/-^ and CS patient fibroblasts after exposure to Illudin S (Fig. 1C), which supports the vector functionality.

**Figure 1.**
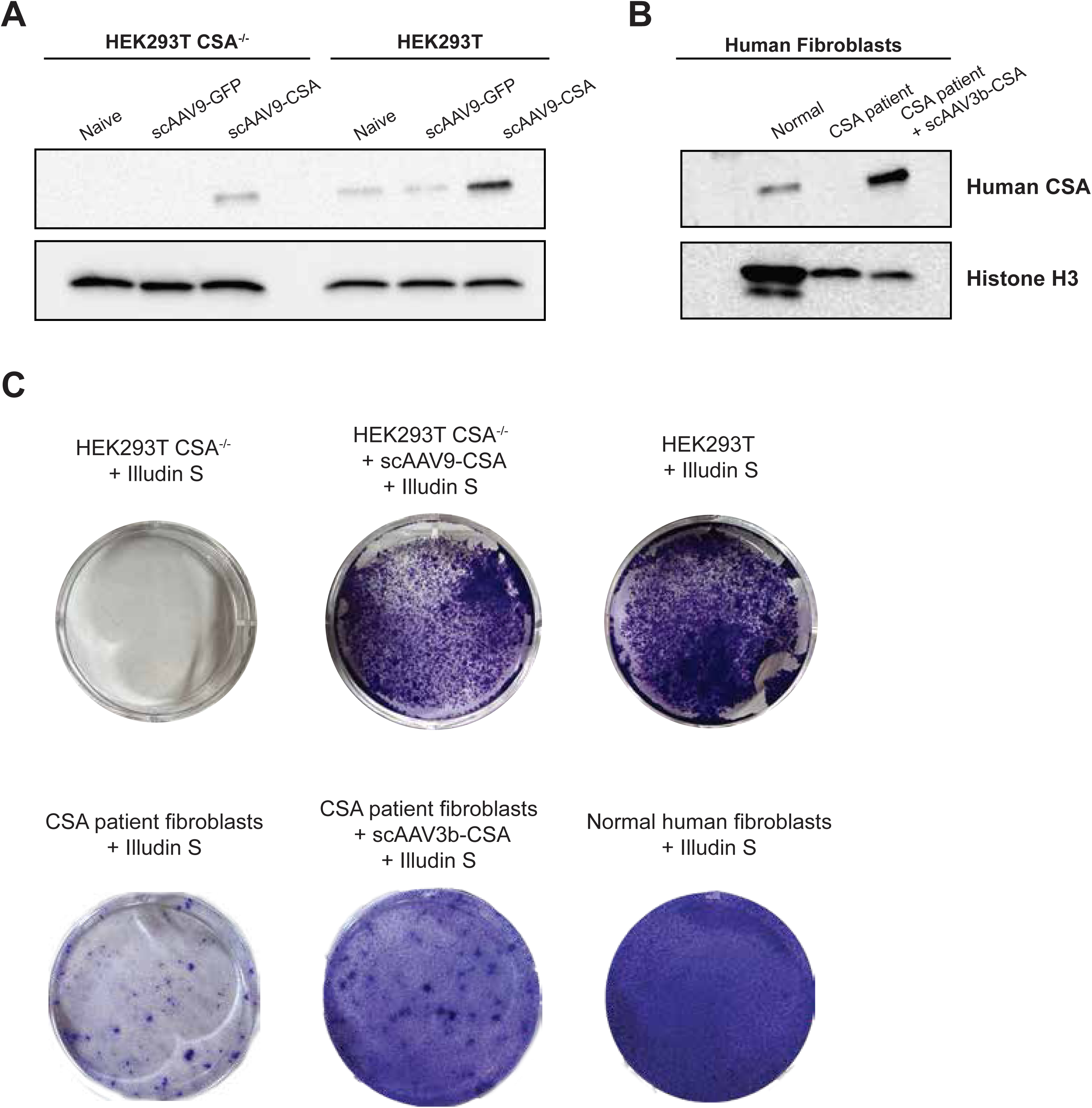
Validation of AAV-CBA-CSA in cell culture. Representative Western blot of CSA and Histone H3 in **(A)** HEK293T and HEK293T-CSA^-/-^ cells and **(B)** human fibroblasts. **(C)** Representative images of Illudin S killing assays. Pictures of individual wells from a 6-well plate are shown.

To assess our vector *in vivo*, we used the very aggressive double knockout for *Csa* and *Xpa* mouse model (*Csa^-/-^; Xpa^-/-^*; referred here as CX mice) (27). Neonatal (P0-P1) bilateral intracerebroventricular (ICV) administration of 1E11vg AAV9-CSA increased significantly (p < 0.0001) the median survival of CX mice from 22 days (PBS injected controls) to 189 days, an 8.5-fold increase (Fig. 2A). At weaning age (21 days), the body weight of AAV treated CX mice was significantly lower than their littermate controls (Fig. 2B). However, their growth rate was comparable to normal littermate controls until ∼ 8 weeks of age (5 weeks post-weaning) when they stopped gaining weight. The inflection point occurred at ∼ 13 weeks of age and body weight decline thereafter until humane endpoint (Fig. 2C). As there were no apparent differences in survival or growth rate between males and females, all animals enrolled in the study were combined into a single group for statistical analyses. Because the treatment was performed at P0-P1, all animals in a single litter were injected independently of genotype. All treated littermates (*Csa^-/-^; Xpa^+/+^* or *Csa^-/-^; Xpa^+/-^*) were euthanized at the same timepoints as AAV9-CSA treated CX mice, with no apparent adverse events.

**Figure 2.**
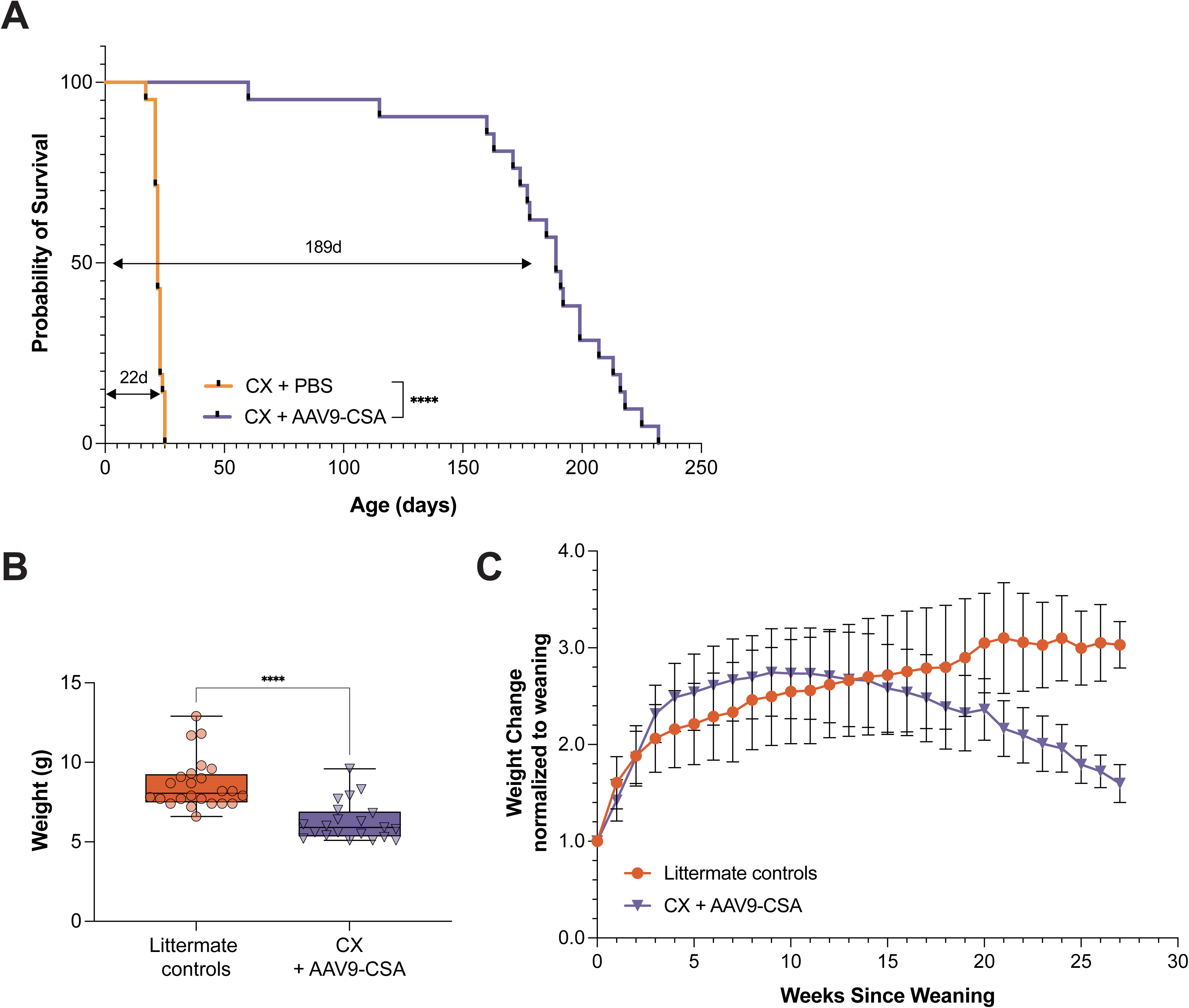
AAV9-CSA treatment increases lifespan and temporary stabilizes growth rate of CX mice. **(A)** Kaplan-Meier survival plot shows 8.5-fold increase in median survival of AAV-treated mice (n=21) compared to untreated mice (n=20). Log-rank (Mantel-Cox) test, p < 0.0001. **(B)** Plot representing the weights at time of weaning (21 days) of AAV-treated mice (n=24, littermate controls + n=21, CX+AAV9-CSA). Data is represented as median and interquartile range. Statistical analysis was performed with two-tailed Mann-Whitney test, p < 0.0001. **(C)** Growth rate of AAV-treated mice, depicting weekly weights normalized to weaning weight of each animal. Data is represented as mean ± SD.

CSA protein expression in brain and heart of AAV-treated and control CX mice at endpoint was evaluated by western blot. We observed variable levels of CSA expression in brain of AAV-treated CX mice (Fig. 3A), which we attribute to injection variability. Interestingly, the differences in CSA expression levels did not correlate with survival. However, CSA expression in heart was more consistent across animals, suggesting equivalent leakage of AAV vector from CSF to the periphery (Fig. 3B). Immunostaining of the brain of AAV-treated CX mice showed widespread distribution of CSA-positive cells throughout the brain (Fig. 3C), with higher expression in cerebral cortex, hippocampus, striatum and in the Purkinje cell layer in the cerebellum, and lower expression in midbrain and upper cortical layers. Our data also shows a predominant neuronal identity of CSA-positive cells in the cortex (Fig. 3D).

**Figure 3.**
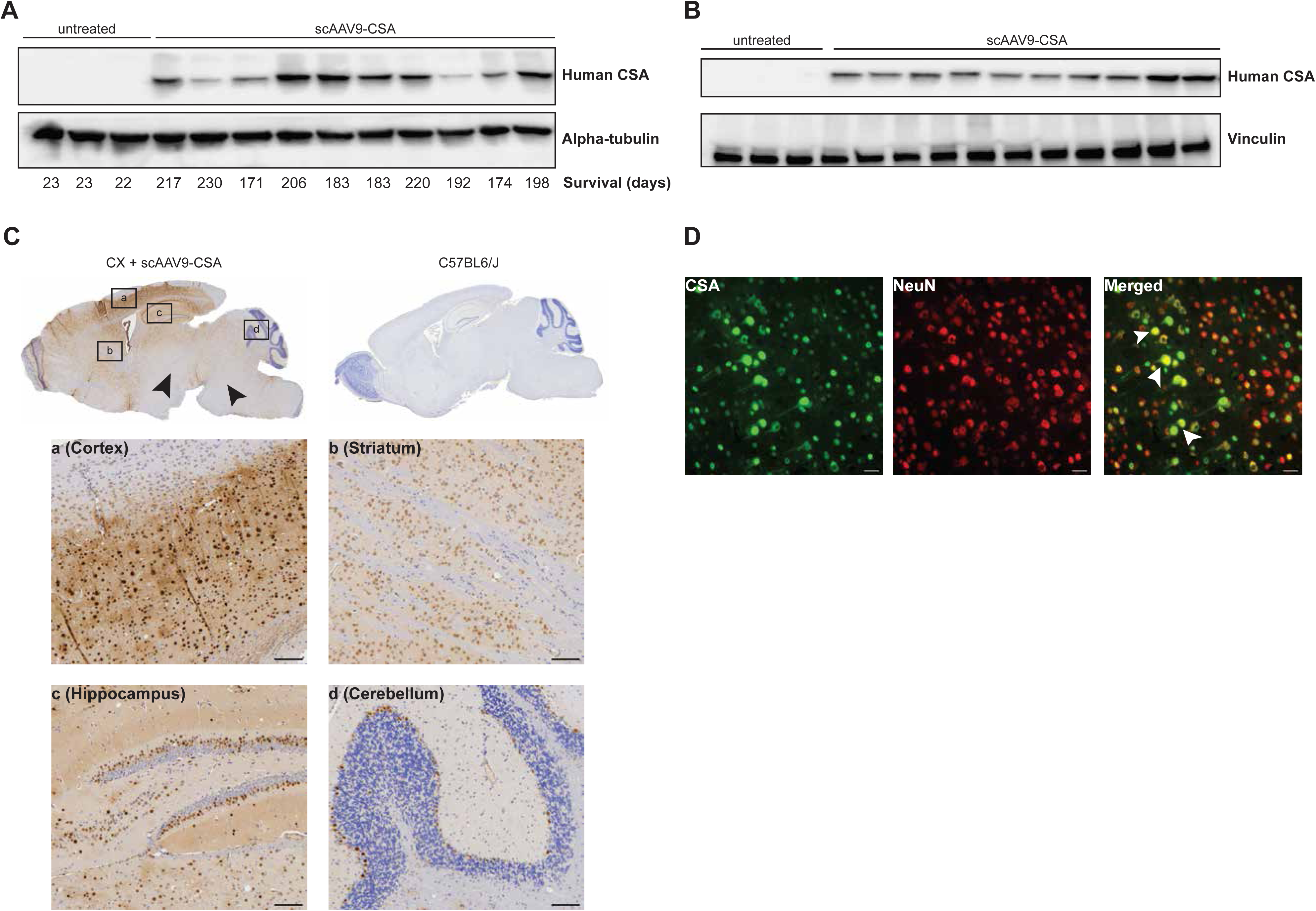
A single neonate ICV injection of AAV9-CSA achieved widespread CSA expression in the brain. CSA protein expression assessed in **(A)** brain and **(B)** heart. Alpha-tubulin or vinculin were used as loading controls. Below the protein blots in **(A)** we show the survival age of each animal in days. **(C)** Representative immunohistochemistry for CSA in CX-treated mice and age-matched C57BL6/J control. Inserts (*a-d*) show high magnification images of different brain regions, indicated in the panel. Black arrows indicate regions of low transduction efficiency. Scale bar, 100 µm. **(D)** Representative immunofluorescence for CSA and NeuN in the cortex of AAV-treated CX mice. Co-localization between CSA and NeuN is shown in yellow (white arrowhead) in the merged panel. Images were acquired as a z-stack and averaged projected. Scale bar, 25 µm.

Hallmarks of CS disease in patients include hypomyelination, calcifications and neuroinflammation in the brain, as well as cranial abnormalities (6, 8, 9, 26, 30). T2-weighted MR imaging of AAV-treated CX mice revealed considerable hypomyelination compared to age matched controls (Fig. 4A). This MRI finding was confirmed by Luxol Fast Blue staining of brain sections, showing lower myelin content in AAV-treated CX mice at humane endpoint, compared to littermates and age matched normal controls (Fig. 4B). Additionally, susceptibility weighted imaging (SWI) revealed magnetic susceptibility artifacts (MSAs) throughout the brain (Fig. 4C-D) consistent with either calcifications or potentially microbleeds from vascular disease (31). However, histological analysis for either bleeds (Prussian blue staining) or calcium (von Kossa staining) proved inconclusive possibly because of small size of the neuropathological alterations and/or the challenge in sampling the exact same brain region. Microbleeds and small calcifications in the brain are known to have an outsized effect on SWI making this technique extremely sensitive for detection of small local events (31).

**Figure 4.**
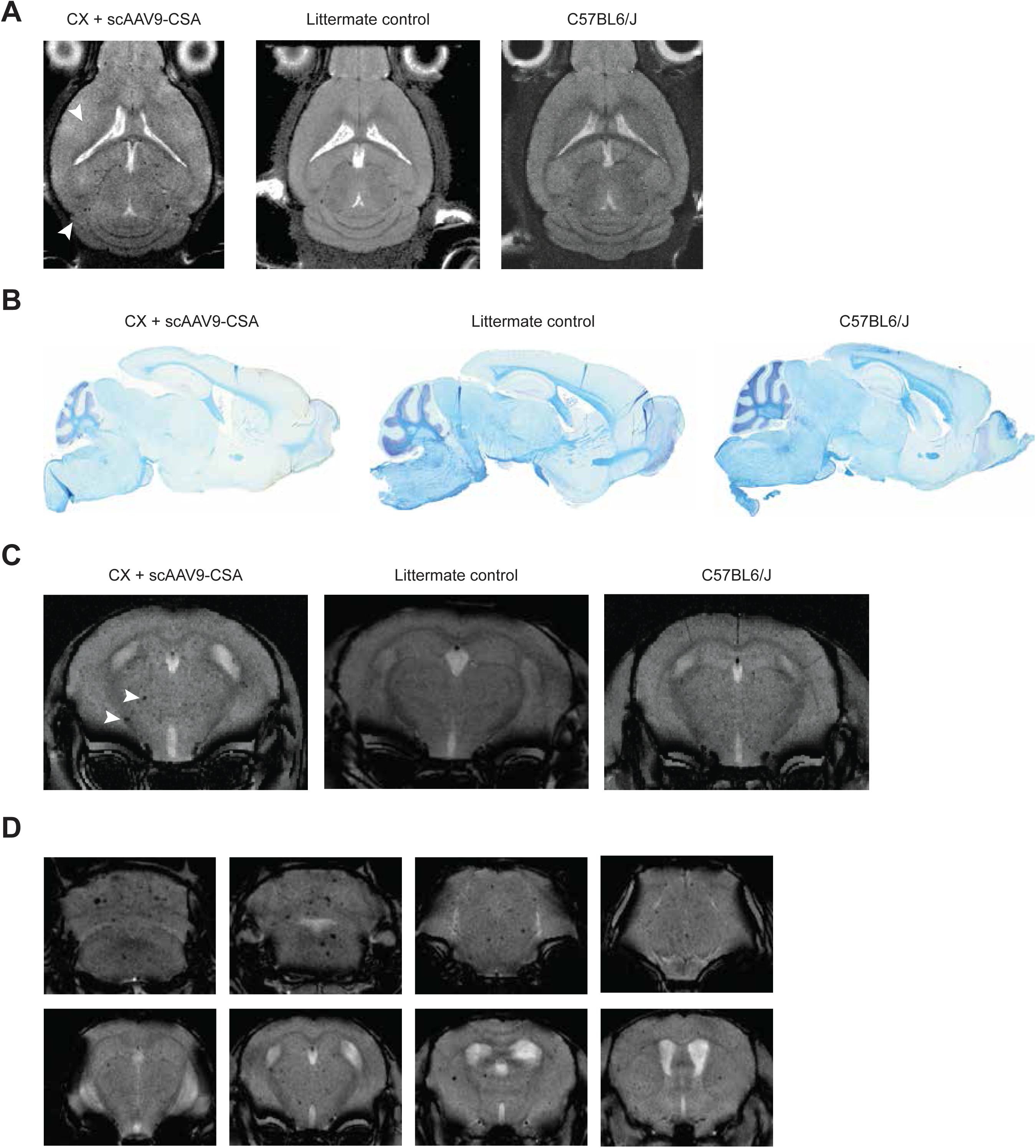
Multimodal brain imaging. **(A)** Representative T2-weighted MRI in AAV-treated CX mice, age-matched littermate controls and C57BL6/J mice shows a lack of gray/white matter differentiation in the CX mice. White arrowheads point at regions where we should see white matter. **(B)** Luxol Fast blue staining of the same animals shows clear differences in myelation status. **(C)** Susceptibility weighted imaging (SWI) shows magnetic susceptibility artifacts (MSA, white arrowheads) indicating possible calcifications or microbleeds. (**D**) SWI images from a single AAV-treated CX mouse single animal showing MSAs in multiple regions throughout the brain.

We also evaluated neuroinflammation by immunofluorescence staining of brain sections with antibodies to markers of reactive astrocytes (GFAP) and activated microglia (Iba1). We found both markers to be significantly elevated in AAV-treated CX mice compared to age-matched WT controls (Fig. 5).

**Figure 5.**
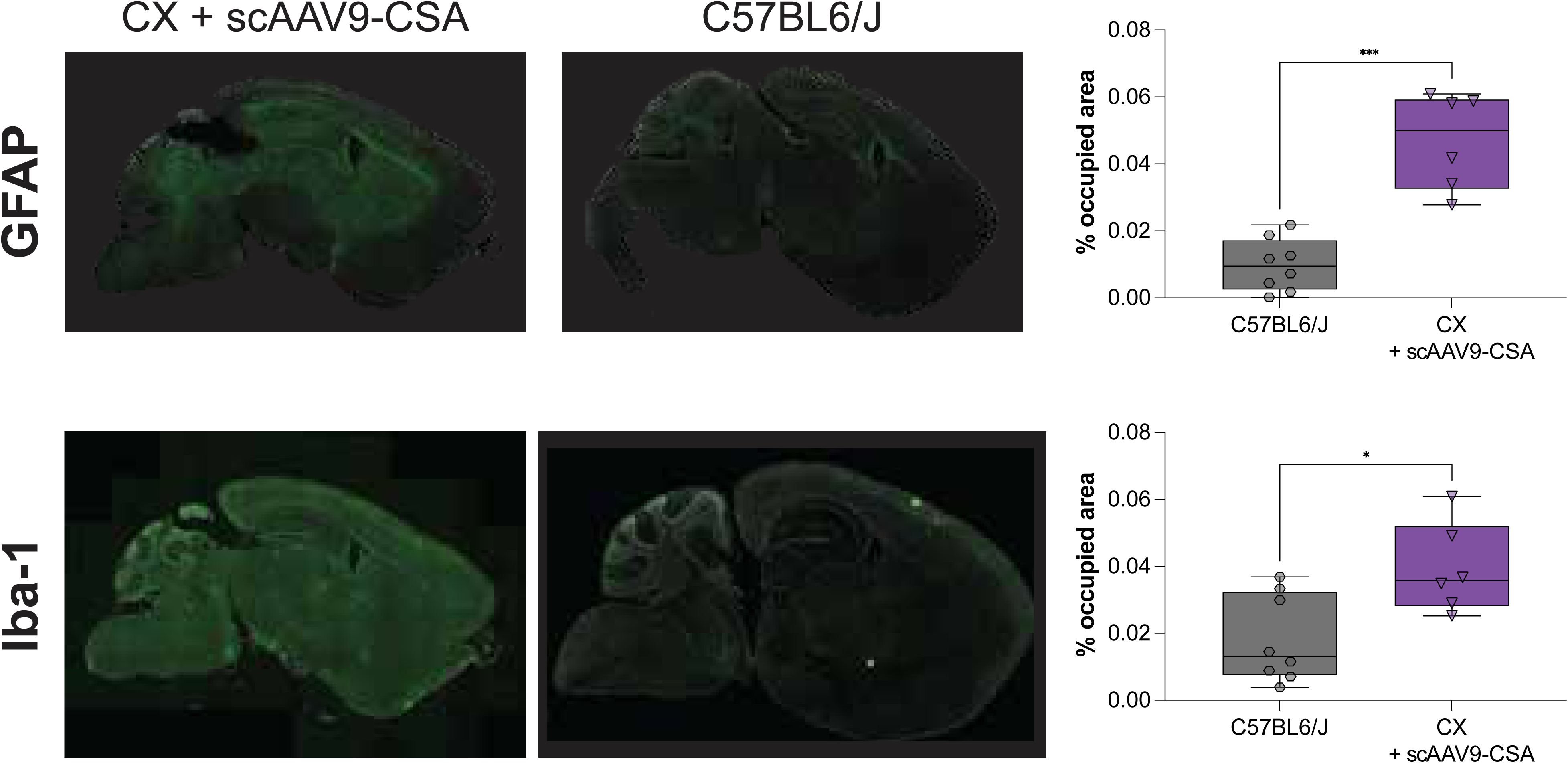
Markers of neuroinflammatory processes. Assessment of the inflammatory markers glial fibrillary protein (Gfap, top panel) or activated microglia (Iba1, bottom panel) in AAV-treated CX mice and age-matched C57BL6/J controls. Staining quantification is represented as median and interquartile range. Statistical analysis was performed with two-tailed unpaired *t*-test with Welch’s correction. For Gfap, p=0.0006; for Iba1, p=0.0136.

Also consistent with disease presentation in CS patients, CT imaging revealed profound kyphosis in untreated CX mice that was not resolved by AAV administration (Sup. Fig. 2A, B). Kyphosis was a criterion for humane endpoint due to its severity. In addition to kyphosis, we also found evidence of skeletal abnormalities in the form of skull surface pitting in AAV-treated CX mice at humane endpoint (Sup. Fig. 2D).

In addition to neurological manifestations, CS patients also present with liver dysfunction as indicated by elevation in serum transaminases and cholestatic enzymes (32). We hypothesized that liver dysfunction may have contributed to growth stalling by ∼ 8 weeks of age and eventual demise of AAV-treated CX mice due to rapid loss of AAV vector genomes in rapidly dividing tissues like the liver in neonatal mice (33). To explore this hypothesis, we performed bulk RNA-seq of livers from AAV-treated CX mice at the humane endpoint and their age-matched littermate controls, who show no signs of disease and have a normal lifespan. In a two-way experimental design with factors for genotype and sex, there were 3,702 genes differentially expressed between genotypes at a cut-off of false discovery rate (FDR) < 0.05 (Fig. 6A, with heatmap of top hits in Fig 6B). There were 167 genes differentially expressed between sexes at the cutoff FDR < 0.05, with the top five hits all consistent with expected sex-related differences: higher levels of four genes on chr Y (*Kdm5d, Uty, Ddx3y, Eif2s3y*) in male vs female mice and lower levels of *Xist* in male vs female mice. There was little evidence of sex-specific expression differences related to *Xpa* genotype, with only 9 genes having FDR < 0.05 in the test for a genotype x sex interaction (Sup. Fig. 3). The differentially expressed genes that were upregulated in KO vs Het mice were over-represented in Molecular Signatures Database hallmark pathways (34) for IL6/JAK/STAT3 signaling, cholesterol homeostasis and hypoxia, while those downregulated were over-represented in metabolic pathways including fatty acid metabolism, bile acid metabolism, oxidative phosphorylation, peroxisome and adipogenesis (Fig. 6C).

**Figure 6.**
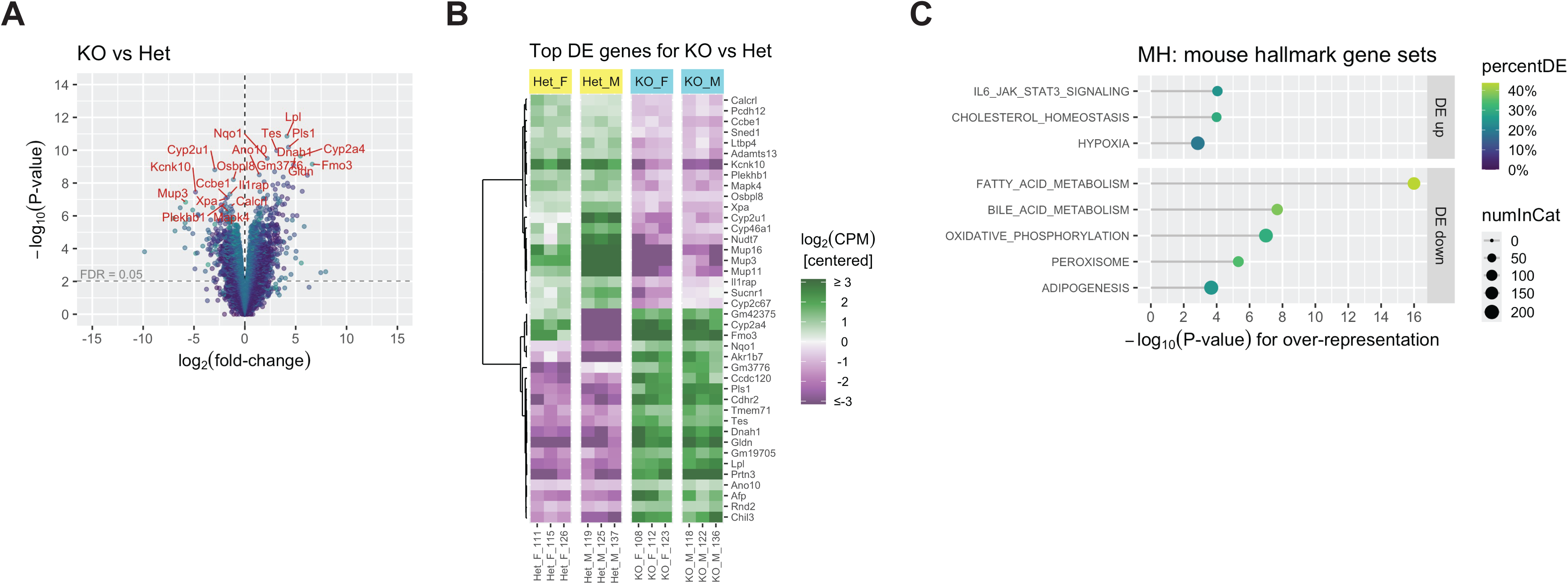
RNA-seq data from mouse liver. (**A**) Volcano plot showing log_2_(fold-change) vs – log_10_(p-value) for differential gene expression in comparison of KO vs Het mice – the main effect of genotype in a model with factors for genotype and sex and their interaction. Color indicates average log_2_(counts-per-million [CPM]) across all samples. Dashed horizontal lines indicate false discovery rate (FDR) = 0.05. (**B**) Heatmap showing the 20 genes in each direction with the smallest p-values in the KO vs Het comparison from (A). Expression levels are on a log_2_ scale, centered to have mean zero for each gene, and capped at +/-3 (i.e., 8-fold above or below mean for the gene). (**C**) Categories from the MSigDB mouse Hallmark gene set collection most significantly enriched among the differentially expressed (DE) genes (genes with FDR < 0.05) from the KO vs Het comparison in (A), separately for DE genes up and down in KO vs Het. Nominal p-values for enrichment are shown. Size of points indicates the number of genes in the category, and color indicates the percent of these genes that were DE in the specified direction.

## DISCUSSION

In this study, we show that administration of AAV9-CSA neonatally to the CSF of an aggressive mouse model of CSA-associated Cockayne Syndrome significantly extended survival, increasing lifespan by 8.5-fold. To our knowledge, this is the first gene therapy for Cockayne Syndrome and with the best therapeutic impact to date, clearly showing the therapeutic potential for CSA gene replacement.

However, it is as important to report therapeutic limitations and AAV-treated CX mice still exhibit a variety of symptoms at endpoint (27, 35), despite the widespread CSA expression. The precise cause of death of AAV-treated CX mice remains unknown. Since we observed no side effects in AAV-treated non-CX littermates, we conclude that the subnormal lifespan of treated CX mice is unrelated to test article related toxicity and more likely related to lack of gene transfer to tissue(s) critical to achieving normal survival.

The natural history of CS in patients includes profound myelination deficits (6, 8). Although there is still debate on whether CS is associated with demyelination or dysmyelination, some studies have characterized it as primarily a hypomyelinating disorder (26). Based on our neuroradiological imaging and histological findings, we were unable to correct myelination with AAV treatment. We believe this is because injection of AAV vectors in P0-P1 mice occurs at a time of rapid oligodendrocyte precursor cells (OPCs) proliferation and the beginning of oligodendrocyte (OL) differentiation only reaches peak levels at P10-P14. In other words, there would be few mature OLs to transduce at the time of AAV injection and AAV vector genomes in transduced OPCs would likely be lost through their rapid proliferation between P0 and P7 and thus not transmitted to the mature OLs that differentiate from OPCs at a later stage of development (36). Additionally, there is evidence that the CBA promoter, which is used in our vector, is not active in OLs (37). Therefore, correcting the hypomyelination component of CS may require using a different promoter that is functional in OLs. Cell-specific promoters derived from the myelin associated glycoprotein (*MAG*) (38) or myelin basic protein (*MBP*) (39) (40) genes are effective for AAV-mediated gene expression in both oligodendrocytes and Schwann cells. These promoters are being used in the development of AAV gene therapies for other diseases affecting nervous system (41). However, restoring CSA expression in only a subset of cell types is unlikely to drive significant therapeutic benefit as this is a ubiquitously expressed protein fundamental for DNA repair through the TC-NER pathway in any cell type.

Other neuropathological findings in CS patients include calcifications, vascular abnormalities (“string vessels”), reactive astrogliosis and microgliosis (30). These are all features we observed in our AAV-treated mice at endpoint. Interestingly, it appears to be some degree of correlation between areas of high astrogliosis (Fig. 5) and areas of poor CSA expression (Fig. 3C), suggesting that we need to improve distribution of CSA expression to the entire brain for full correction of the neurological aspects of the disease.

Besides the neurological findings, CS patients also present with elevated liver transaminases, lipodystrophy, cholestasis and other signs of liver dysfunction (6, 8). The transcriptomic changes documented in the liver of endpoint AAV-treated CS mice are consistent with these findings in patients. Upregulation of genes represented in the IL6/JAK/STAT pathway are indicative of an ongoing inflammatory process in liver which is normally accompanied by elevation in serum transaminases. Interestingly downregulated genes were in fatty acid and bile acid metabolism pathways as well as adipogenesis pathways which may explain the metabolic alterations in CS patients. Therefore, it is our hypothesis that metabolic dysfunction was a major determinant of the shorter-than-normal lifespan of AAV-treated CX mice. It is well established that mice treated neonatally with AAV vectors lose liver genomes overtime due to hepatocellular turnover resulting in loss of therapeutic effect (33, 42). This is in line with our finding of adipogenesis dysregulation, and failure of AAV-treated CX mice to sustain weight gain. This again parallels the lipodystrophy (reduced subcutaneous fat) documented in CS patients and it is likely caused by liver dysfunction (6, 8, 32, 43). However, hepatic turnover is an experimental limitation of studies in neonatal animals and should not be a major cause of concern for clinical translation.

Modelling the complexity of CS in small rodents has proven to be a challenging task, as animal models often fail to reproduce disease phenotypes (25). Our CX model is a double mutant for *Csa* and *Xpa*, with both genes involved in the TC-NER pathway. While this is an aggressive model that allows for faster testing of therapies, presently we are unable to exclude the possible contribution of *Xpa* mutations to the lingering phenotypes we observe in AAV-treated CX mice, even though the *Xpa^-/-^*mouse model has no overt phenotype (44, 45). Furthermore, CSA is part of a complex protein machinery that requires coordinated interactions of multiple proteins for appropriate responses to stress (21, 46). We cannot discard the possibility that expressing the human protein in mouse cells, may not allow for the proper assembly of repair complexes and could be the cause for the incomplete correction of phenotypes in the CX mice.

Additionally, CSA is ubiquitously expressed (47), and therefore a successful gene therapy needs to transduce virtually every cell to achieve a full correction. Although we observed widespread transduction in the cortex, deficiencies were noted in deep brain structures (Fig. 3). Therefore, further research is still warranted to develop better therapies for CS. The profound neurological and peripheral symptoms in CS indicate that a transformative AAV gene therapy will require new capsids that target the CNS at higher efficiency than AAV9 and retain its broad peripheral tissue tropism. The new generation of AAV capsids identified by *in vivo* or *in vitro* screening of capsid libraries are orders of magnitude more potent than AAV9 for CNS gene transfer (48-51) (52-54). Importantly these new AAV9 variants largely retain the broad peripheral tissue tropism with the added advantage that liver tropism is often tuned down but not altogether eliminated. This is an important consideration when selecting an AAV capsid for development of a gene therapy for neurological diseases like CS with CNS and peripheral pathology as exemplified by the apparent liver disease in CX mice.

The second component of a successful gene therapy for broadly expressed genes such as CSA is to use promoters with broad functionality expressing the therapeutic protein at normal or nearly normal physiological levels. Development of transgene expression cassettes with broad functionality can be achieve using combinations of cell specific promoters (37) or new promoters with endogenous-like expression (42). In recent years bioinformatic analysis of single cell transcriptomic and epigenomic data from mouse and human has been used to developed new enhancer elements with exquisite specificity for different cell populations in CNS (55-58) (59). This same approach may be also applicable to develop small gene-specific promoters capable of expressing the respective therapeutic protein close to physiological levels, or AAV-compatible promoters derived from housekeeping genes likely to function in most cells. The combination of potent new AAV capsids for CNS gene delivery with the emerging principles of promoter engineering based on single cell multiomics will be the basis for development of a second generation AAV gene therapy with the potential to achieve transformative therapeutic outcomes for CS patients.

In conclusion, our data supports the notion that our AAV9-CSA vector is a promising therapeutic approach for the treatment of CSA-linked Cockayne syndrome, based on extension of lifespan alone. However, some questions remain unanswered and pave the way for further research into better vectors that could translate into clinical trials and substantially impact patients’ lives.

## METHODS

### AAV vector design and preparation

The self-complementary AAV vector used in these studies carries a transgene cassette with the wild type human *ERCC8/CSA* reference cDNA (NM_000082.4), which encodes the longest isoform of CSA, driven by a CBA promoter composed of the cytomegalovirus immediate-early gene enhancer upstream of the chicken beta-actin (CBA) promoter and followed by an SV40 intron, and it uses a BGH (bovine growth hormone) polyA signal for polyadenylation. The transgene cassette is flanked by AAV2 inverted terminal repeats (ITR). The ITR near the CMV enhancer carries a deletion of the terminal resolution (TR) sequence for production of self-complementary AAV genomes during packaging.

AAV9 vector was produced, purified and its titer determined as previously described (60). Throughout this study, several independent batches of vector were required. All titers were normalized to the value of the first batch to ensure consistency of dosing.

### Cell lines and *in vitro* assays

We generated CSA deficient HEK293T cells (HEK293T-CSA^-/-^) by using CRISPR/Cas9 to disrupt exons 2 and 7 of the *ERCC8* gene. The efficacy of disruption was validated by Western blot of cell lysates, as described below (Sup. Fig. 1). HEK293T and HEK293T-CSA^-/-^ cells were cultured in DMEM (12430-047, Gibco) with 10% FBS (Sigma-Aldrich), 1% GlutaMAX-I (Gibco) and 100 U/mL penicillin/streptomycin (Gibco). Normal human fibroblasts (AG08498) and CSA patient fibroblasts (GM28257) were acquired from the Coriell Institute for Medical Research. Both cell lines were cultured in EMEM (Gibco) supplemented with 15% FBS (Sigma-Aldrich), 1% GlutaMAX-I (Gibco), 1% (v/v) non-essential amino acids (Gibco) and 100 U/mL penicillin/streptomycin (Gibco). All cell lines were routinely propagated as monolayers at 37°C in 5% CO_2_ humidified incubators.

#### Assessing CSA protein expression

Cells were plated in 12-well plates at a density of 300,000 cells per well (HEK293T) or in 6-well plates at 1,000,000 cells per well (fibroblasts), and 24 h later they were infected with AAV-CSA or AAV-eGFP (as control) at an MOI of 100,000 vg/cell. Experiments in HEK293T were performed with AAV9 vectors, whereas experiments in fibroblasts used AAV3b vectors. At 72 h post-infection, cells were assessed for eGFP expression by fluorescence microscopy, lysed in RIPA buffer (50 mM Tris pH 7.4, 140 mM NaCl, 5 mM EDTA, 0.1% SDS, 0.5% sodium deoxycholate, 1% NP-40) with protease inhibitors (cOmplete Mini, EDTA-free, Sigma Aldrich) and processed for Western blot analysis as described below.

#### Illudin S assays

All cell lines were plated in 12-well plates (300,000 cells per well) and infected as above. At 36 h post-infection, cells were trypsinized and replated in 6-well plates at a density of 100,000 cells per well. Twenty-four hours later, Illudin S (17451, Cayman Chemical) was added to the cells at a concentration of 4 ng/mL (HEK293T) or 2 ng/mL (fibroblasts). Cells were fixed in formalin and stained with Crystal Violet at 72 hrs after adding Illudin S.

### Animal procedures

*Csa^-/-^;Xpa^+/-^* embryos were kindly donated by Drs. Michael MacArthur and Sarah Mitchell (Jay Mitchell group, ETH Zurich, Switzerland) and recovered at the UMass Transgenic Mouse Core. Seven-month-old C57BL/6J mice were purchased from The Jackson Laboratory (stock no. 00064). Animals were maintained at 21 ± 1 °C under a 12 h light/dark cycle with water provided *ad libitum*. Breeding pairs were fed a 10% kcal fat diet (D12450Bi, Research Diets), while all study animals after weaning were provided with regular irradiated laboratory chow.

#### Sex as a biological variable

All our study groups include male and female animals. Similar findings are reported for both sexes, with no differences in survival or growth rate or transcriptome analysis, between males and females. Therefore, both sexes were combined into a single group for statistical analyses.

#### Intracerebroventricular (icv) delivery

P0-P1 pups were anesthetized using isoflurane (drop method) and injected with 1 x 10^11^ vg of AAV9-CSA freehand in the lateral ventricles (2 µL + 2 µL) using a 10 µL glass Hamilton syringe fitted with a 32 G beveled needle. Animals were returned to home cage with mom and monitored until full recovery.

All animals in the same litter were injected regardless of genotype. Mice were genotyped at weaning. CX mice (*Csa^-/-^;Xpa^-/-^*) were followed until humane endpoint, defined by severe hunched posture and body tremors, at which point they were euthanized with an overdose of ketamine (375 mg/kg) and xylazine (37.5 mg/kg) and multiple organs were collected and bisected, with one portion immediately frozen (for molecular analysis) and the other fixed in 10% neutral buffered formalin and embedded for paraffin processing. All non-CX littermates (*Csa^-/-^;Xpa^+/-^* or *Csa^-/-^;Xpa^+/+^*) were used as controls and euthanized in the same manner.

### Radiological imaging

A subset of AAV-treated CX mice (n=5) were subjected to radiological imaging at endpoint before euthanasia, as were age-matched littermate controls (n=8) and normal C57BL6/J mice (n=6). Animals were subjected to both 7T MRI (Bruker biospec 70/30, Billerica, MA) and microCT (VectorCT, MI labs, Houten, Netherlands). A standard imaging protocol was used for each animal including qualitative imaging in the form of coronal T2 weighted (TR/TE: 2500/33 ms, FoV: 15x15 mm, matrix: 256x256, Thickness: 0.5 mm, NSA: 4) and susceptibility weighted imaging (SWI) (TR/TE: 425/10 ms, FA: 30°, FoV: 20x15 mm, matrix: 280x210, Thickness: 0.5 mm, NSA: 4). Quantitative diffusion imaging in the form of diffusion tensor imaging (DTI) (TR/TE: 2500/21 ms, FoV: 18x15 mm, matrix: 108x90, Thickness: 0.8mm, NSA: 2, directions: 30) was also acquired. After the MRI protocol was completed, the animals were transferred to the microCT system for whole body computed tomography (20 µm isotropic resolution) to assess bone health and qualitative spinal curvature. For both imaging modalities animals were anesthetized with isoflurane gas (1-3%) and allowed to spontaneously breath air with 1.5% isoflurane.

### Histology and Image processing

#### Immunohistochemistry

Four-µm-thick sections were deparaffinized in Xylene and hydrated in a descendent alcohol series. Heat induced antigen retrieval was performed using a microwave and sodium citrate buffer (10 mM sodium citrate, pH 6.0). Tissue slides were incubated with blocking buffer containing 5% fetal bovine serum and 5% normal goat serum in phosphate-buffered saline for 1 h at room temperature and then incubated overnight at 4°C with primary antibodies diluted in blocking buffer. Primary antibody used was rabbit recombinant monoclonal anti-ERCC8 antibody [EPR9237] (1:100, Abcam). Endogenous peroxidase activity was quenched with 3% hydrogen peroxide in phosphate-buffered saline for 20 min. Antigen visualization was performed using Vectastain Elite ABC Reagent and DAB Substrate Kit (both from Vector Labs), according to the manufacturer’s instructions. Sections were counterstained with hematoxylin, dehydrated in an ascending alcohol series and coverslipped using Permount (Fisher Scientific). Sections were visualized under a Leica Thunder Imager DMi8 microscope equipped with a DMC4500 digital camera.

#### Immunofluorescence

Four-µm-thick sections were deparaffinated in Xylene and hydrated in a descendent alcohol series. Heat induced antigen retrieval was performed using a microwave and sodium citrate buffer (10 mM sodium citrate, pH 6.0). Tissue slides were incubated with blocking buffer containing 5% fetal bovine serum, 5% normal donkey serum and 0.3 M glycine in phosphate-buffered saline for 1 h at room temperature. Primary antibodies were diluted in blocking buffer without glycine and then incubated overnight at 4°C. Primary antibodies used were rabbit recombinant monoclonal anti-ERCC8 [EPR9237] (1:100, Abcam), rat monoclonal anti-NeuN [EPR12763] (1:100, Abcam); rabbit polyclonal anti-glial fibrillary acidic protein (1:1,000, Agilent); rabbit polyclonal anti-Iba1 (1:1,000, Wako). Sections were then washed and incubated with secondary antibodies AlexaFluor-555 anti-rabbit IgG or AlexaFluor-647 anti-rabbit IgG (1:1,000; Invitrogen Molecular Probes) for 1 h at room temperature. Slides were mounted in Vectashield containing DAPI (Vector Labs) and visualized under a Leica Thunder Imager DMi8 microscope equipped with a Leica K5 digital camera and using the same exposure conditions across groups.

For both staining techniques, parallel sections were incubated with blocking buffer in lieu of primary antibody as staining controls, to assess antibody specificity. Staining was absent under these conditions.

#### Image processing

All images were processed using either Fiji software (61) or in-house developed MATLAB algorithms, and changes to brightness or contrast or gamma were kept consistent across groups. Tissue sections immunostained for GFAP and IBA-1 expression were analyzed using the same algorithm. The image quantification algorithm is a two-step automated process: First the images were processed through a Gaussian filter to smooth the noise. These filtered images were then thresholded and a marching squares algorithm (62) applied to segment the outer boundary of the brain. The area inside this segmentation defines the brain area. Once the brain area was defined the original unfiltered image was subdivided into smaller regions for parallel processing. Each subsection was thresholded to detect only the stained cells (GFAP or IBA-1) using an adaptive method that accounts for the local background intensity variations. The total area of all the stained cells was then summed across all subsections. The final result is presented as the percent area of the brain that is occupied by the stained cells.

### Western Blot

For protein expression analysis, tissues were lysed in RIPA buffer containing protease inhibitors, as above. Total protein in the lysates was determined by Bradford assay (Bio-Rad) using serial dilutions of bovine serum albumin as protein standard; 40 µg of total protein was separated in a 4-20% polyacrylamide SDS-PAGE gel (Bio-Rad) and then transferred to 0.2 µm pore nitrocellulose membrane (Amersham, GE Healthcare). Primary antibodies were rabbit recombinant monoclonal anti-ERCC8 [EPR9237] (1:100, Abcam), mouse monoclonal anti-α-tubulin (1:1,000; T6199, Sigma-Aldrich) or rabbit monoclonal anti-vinculin (1:1,000; ab129002, Abcam) for normalization. Detection was performed by chemiluminescence using Clarity Western ECL Substrate (Bio-Rad) and images were acquired using a ChemiDoc system (Bio-Rad).

### RNAseq

Total RNA was isolated using TriZol Reagent (Invitrogen) according to the manufacturer instructions. Libraries were prepared with ribosomal-RNA depletion (TruSeq Stranded Total RNA Ribo-Zero H/M/R Gold kit; Illumina) and sequenced on the NovaSeq X platform by Psomagen with 50-70M 2x151 paired-end reads per sample (with 12 samples total: livers from 3 mice per sex per genotype).

The nf-core/rnaseq (v3.14.0)(63) pipeline from the nf-core collection (64) of NextFlow workflows(65) was used for data QC and preprocessing, for mapping reads with the STAR aligner (2.7.9a) (66) to the mouse GRCm39/mm39 reference genome with GENCODE M34/Ensembl 111 annotations, and for computing estimated read counts per gene per sample with Salmon (1.10.1) (67). The R (v.4.3.2) package edgeR (v.4.0.16) (68) was used to perform quasi-likelihood tests (69) for differential gene expression in a generalized linear model with factors for genotype (KO = *Csa^-/-^;Xpa^+/+^* and Het = *Csa^-/-^;Xpa^+/-^*), sex (M and F), and their interaction (with non-default parameters prior.count=2 and robust=T for glmQLFit). Genes that did not have at least 5 reads in at least 3 samples (after adjusting for differences in library sizes while keeping median library size fixed) were filtered out prior to edgeR analysis. FDR (70) was used to control for multiple hypothesis testing. The R package goseq (v.1.54.0) was used for gene set analysis, testing for enrichment of categories from the Molecular Signatures Database (MSigDB v2023.2.Mm) (71) among differentially expressed genes (FDR < 0.05), using log2(CPM) [CPM = counts per million] as a covariate to adjust for potential biases due to gene expression level (72).

### Statistical analysis

All statistical analysis were performed using GraphPad Prism v10.4 for macOS, aside from RNA-seq analysis described above. Statistical tests on survival experiments were performed using Kaplan-Meyer analysis and Log-rank (Mantel-Cox) tests. For all other tests, data is plotted as median and interquartile range. Prior to statistical analysis, normality testing was performed. Comparisons between groups that did not fail tests for normality, were analyzed with parametric tests. Otherwise, non-parametric tests were used. Details on specific analysis are detailed in the figure legends.

### Study Approval

All animal procedures were performed as approved by the Institutional Animal Care and Use Committee at UMass Chan Medical School (PROT202100108).

## Data Availability

All raw data and MATLAB scripts are available upon request.

## AUTHOR CONTRIBUTIONS

ARB and MSE conceived the study and designed the experimental plan. ACS, MKW, TM, KOH performed animal experiments. ACS, WSC, MKW, CMS processed samples and collected data. ARB analyzed data. ODK analyzed RNA-seq data. RMK performed radiological imaging and staining quantification. ARB and MSE wrote manuscript. MSE supervised the project.

## ACKNOWLEDGEMENTS

We thank Heather Gray-Edwards for invaluable edits on the manuscript draft. We also thank the Radio Labeling Small Animal Imaging Core Facility and Advance MRI Center at UMass Chan Medical School for their support in the radiological imaging studies. We are deeply grateful to Sarah Mitchell and Michael MacArthur and the entire Jay Mitchell group at ETH Zurich, Switzerland for kindly providing us with the last *Csa^-/-^;Xpa^+/-^* mouse embryos.

This study was supported by the Riaan Research Initiative Foundation.

## SUPPLEMENTARY FIGURES

**Supplementary Figure 1.**
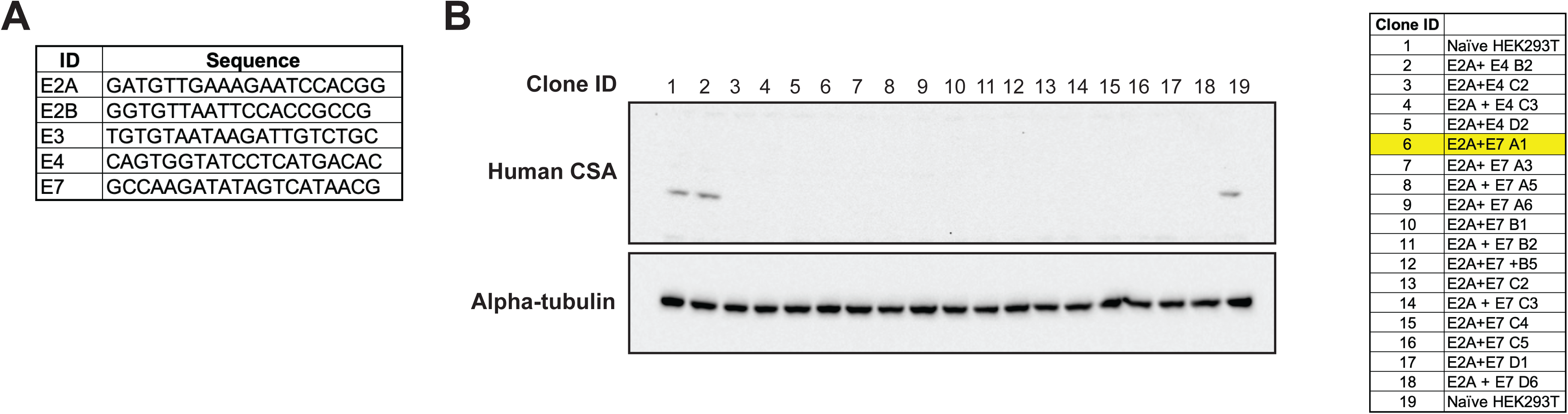
Generation of HEK293T *Csa^-/-^* cell line. We generated a CSA KO cell line by co-transfection of HEK293T cells with plasmids expressing gRNAs, Cas9 and puromycin resistance gene. Sequence of all gRNA used in (**A**), with indication of the location in the ID name (for example, E3 means it targets exon 3). Different clones were then selected in media with 1 mg/mL puromycin (Invivogen) and tested for CSA expression by WB (**B**). Table shows all the clones and gRNA combinations tested. Clone number 6 (highlighted in yellow) was selected as the CSA KO line for our studies. Of note, only one of 17 clones tested still expressed CSA (clone #2).

**Supplementary Figure 2.**
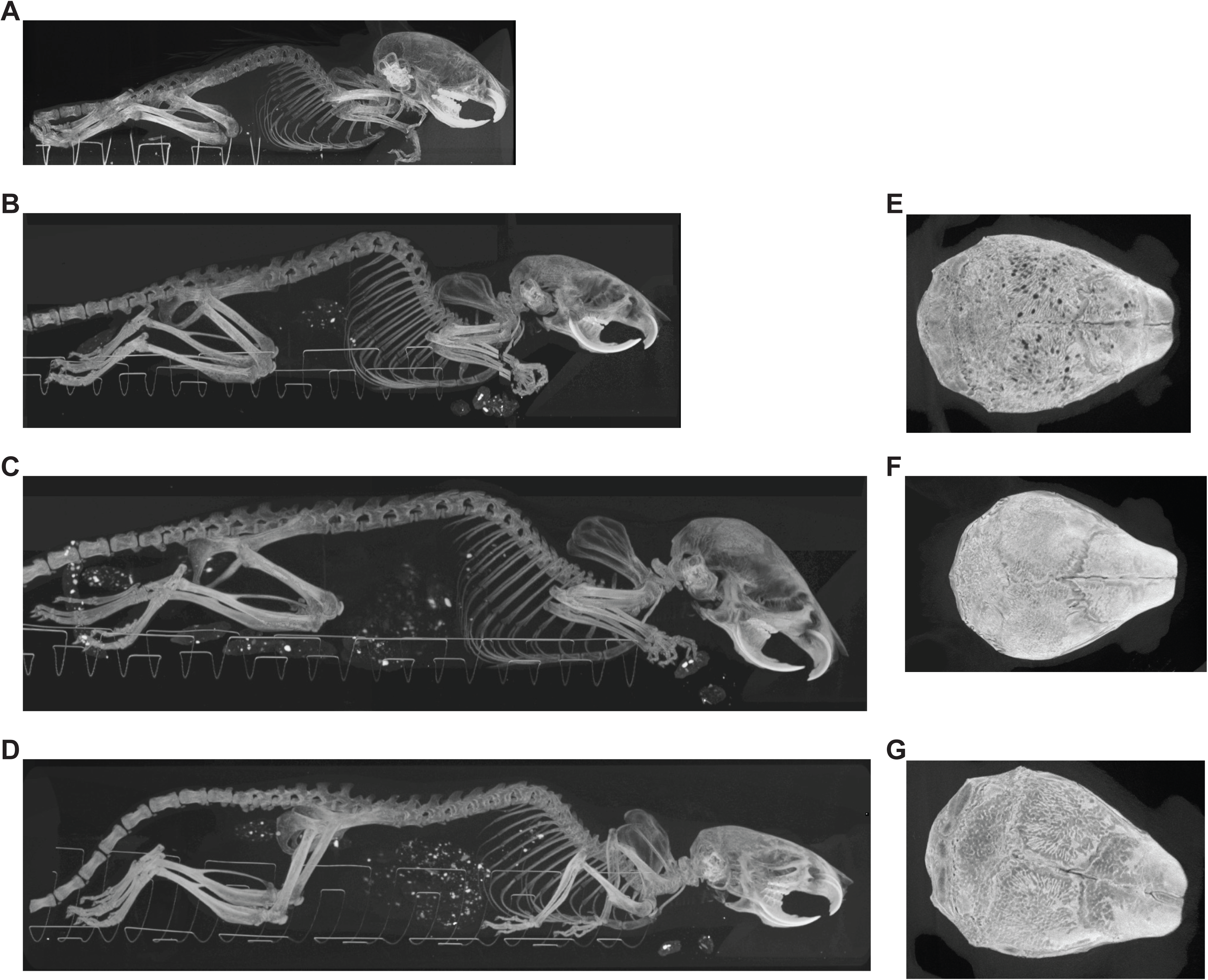
CT imaging. Representative maximum intensity projection (MIP) reconstruction of whole body microCT for (**A**) untreated CX mouse at endpoint (22 days old); (**B**) AAV-treated CX mouse at endpoint (approximately 7 months old); (**C**) age matched littermate and (**D**) C57BL6/J control. Note the severe kyphosis in (A) and (B). Skull MIP revealed considerably surface pitting in (**E**) AAV-treated CX mice at endpoint compared to (**F**) age matched littermates and (**G**) C57BL6/J controls.

**Supplementary Figure 3.**
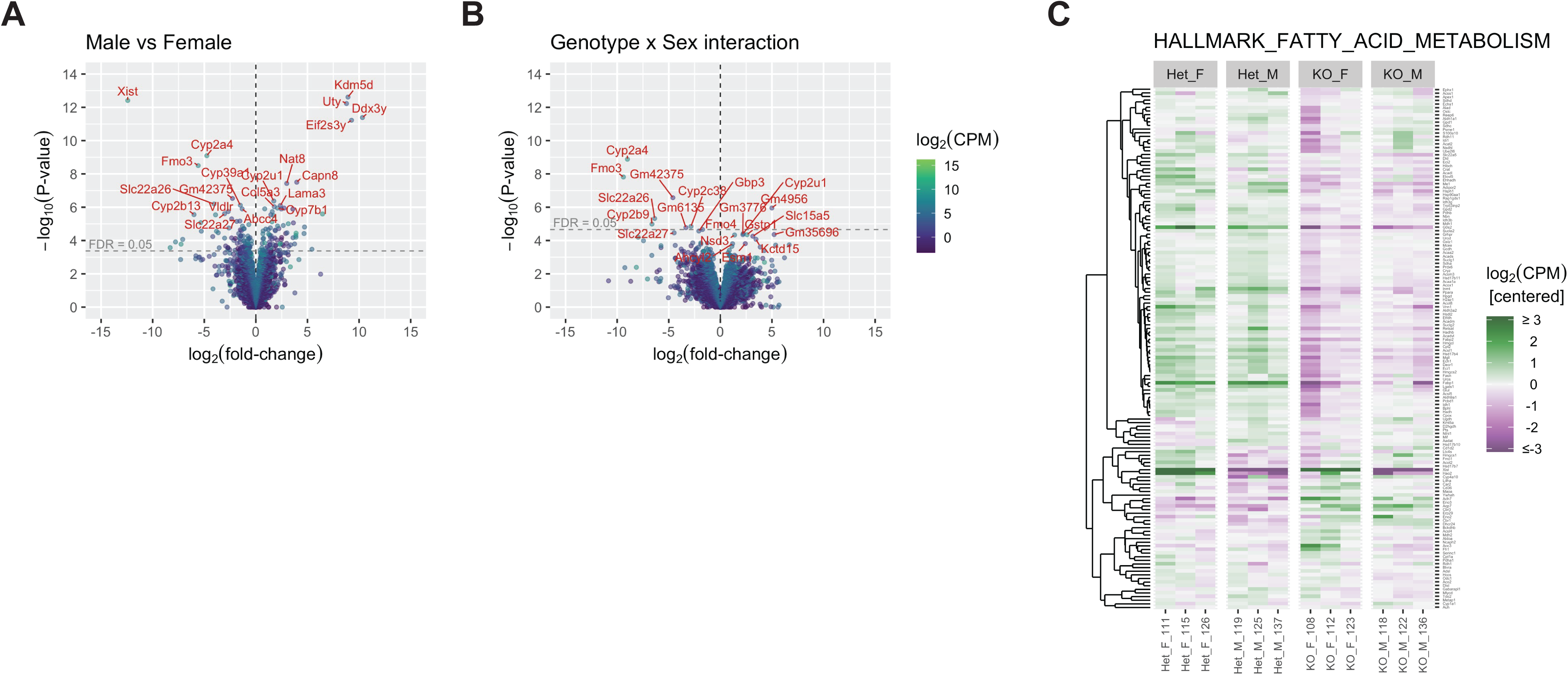
RNA-seq data from mouse liver. Volcano plots showing log_2_(fold-change) vs –log_10_(p-value) for differential gene expression in (**A**) male vs female mice, and (**B**) the difference in KO vs Het for male vs female mice (i.e., Xpa genotype x sex interaction). Dashed horizontal lines indicate false discovery rate (FDR) = 0.05; at this cutoff there were 167 in (A), and 9 in (B), out of 19440 genes tested after filtering out genes with very low expression. Note that the top hits for (A) are *Xist* (down in male vs female) and four genes on the Y-chromosome (up in male vs female), consistent with expected sex-related differences, and that despite widespread expression differences associated with *Xpa* genotype there is little evidence for sex-specificity of these differences in (B). (**C**) Heatmap for all the genes in the “hallmark fatty acid metabolism” gene set, the most sig hit from pathway analysis. Expression levels are on a log_2_ scale, centered to have mean zero for each gene, and capped at +/-3 (i.e., 8-fold above or below mean for the gene).

